# Imaging crossing fibers in mouse, pig, monkey, and human brain using small-angle X-ray scattering

**DOI:** 10.1101/2022.09.30.510198

**Authors:** Marios Georgiadis, Miriam Menzel, Jan A Reuter, Donald Born, Sophie Kovacevich, Dario Alvarez, Zirui Gao, Manuel Guizar-Sicairos, Thomas M Weiss, Markus Axer, Ivan Rajkovic, Michael M Zeineh

## Abstract

Myelinated axons (nerve fibers) efficiently transmit signals throughout the brain via action potentials. Multiple methods that are sensitive to axon orientations, from microscopy to magnetic resonance imaging, aim to reconstruct the brain’s structural connectome. As billions of nerve fibers traverse the brain with various possible geometries at each point, resolving fiber crossings is necessary to generate accurate structural connectivity maps. However, doing so with specificity is a challenging task because signals originating from oriented fibers can be influenced by brain (micro)structures unrelated to myelinated axons.

X-ray scattering can specifically probe myelinated axons due to the periodicity of the myelin sheath, which yields distinct peaks in the scattering pattern. Here, we show that small-angle X-ray scattering (SAXS) can be used to detect myelinated, axon-specific fiber crossings. We first demonstrate the capability using strips of human corpus callosum to create artificial double- and triple-crossing fiber geometries, and we then apply the method in mouse, pig, vervet monkey, and human brains. Given its specificity, capability of 3-dimensional sampling and high resolution, SAXS can serve as a ground truth for validating MRI as well as microscopy-based methods.

**Statement of Significance:** To study how the nerve fibers in our brain are interconnected, scientists need to visualize their trajectories, which often cross one another. Here, we show the unique capacity of small-angle X-ray scattering (SAXS) to study these fiber crossings without use of labelling, taking advantage of SAXS’s specificity to myelin - the insulating sheath that is wrapped around nerve fibers. We use SAXS to detect double and triple crossing fibers and unveil intricate crossings in mouse, pig, vervet monkey, and human brains. This non-destructive method can uncover complex fiber trajectories and validate other less specific imaging methods (e.g., MRI or microscopy), towards accurate mapping of neuronal connectivity in the animal and human brain.

**Graphical Abstract:** 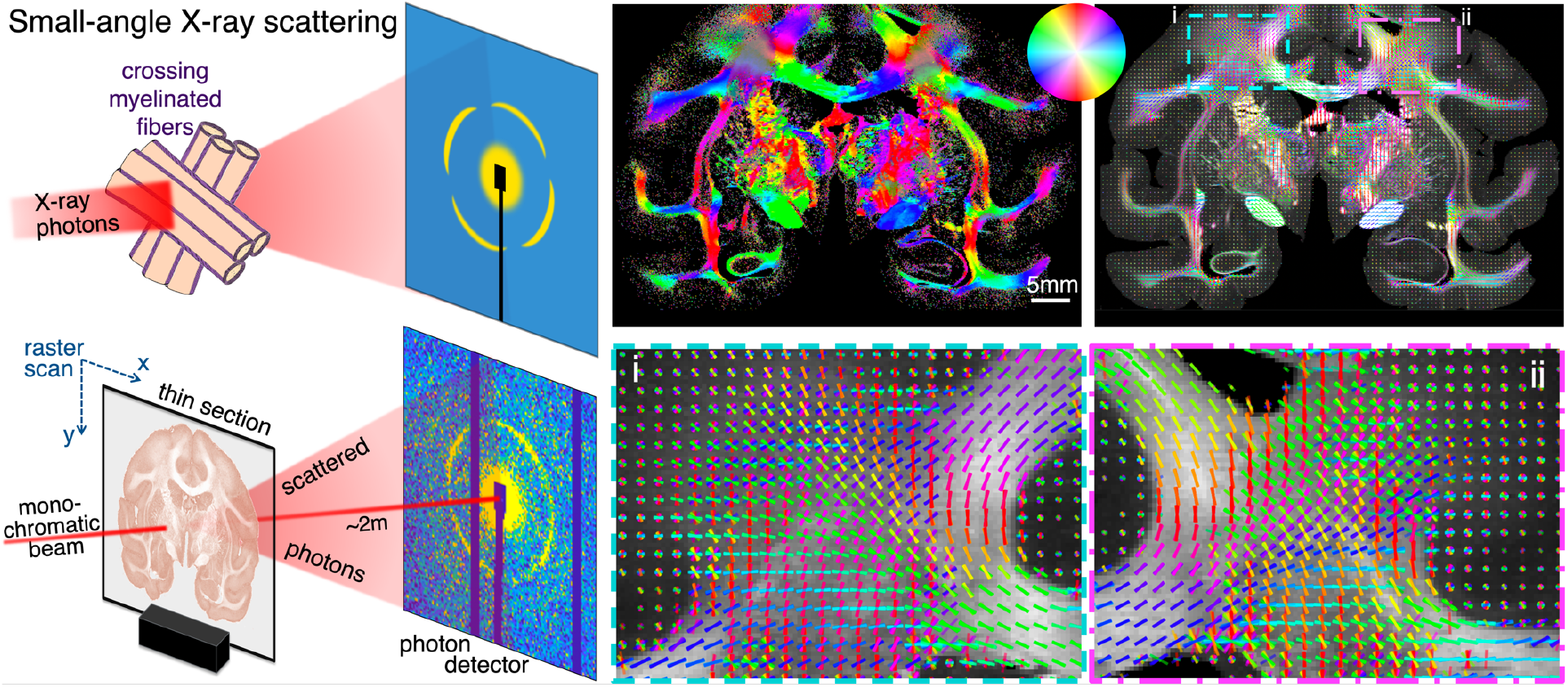

## 1 Introduction

Signals in the brain are transmitted within neurons via action potentials. These are typically initiated in the soma of neurons, and travel through neuronal axons (nerve fibers) until they reach the synaptic clefts where biochemical mechanisms further transmit the signals to the next cell. A major evolutionary step in the signal transmission in the brain came with myelination: oligodendrocytes, an abundant type of central nervous system glial cells found in vertebrates, form processes that “wrap around” neuronal axons (Schwann cells do the same in the peripheral nervous system). The resulting layered structure around axons is called myelin (with a layer periodicity of ∼15-20nm) and is an essential component of our brain, constituting ∼35% of its dry weight [1].

In the beginning of the 21^st^ century, the “brain connectome” was conceived [2], [3], with the goal of mapping neuronal connections across the animal and human brain. Given that there are >50 billion neurons in a human brain [4], with more than 100,000 km total length of myelinated fibers [5], each of diameter from 0.1 to 100 micrometers [6] (with most axons being around 1μm in diameter), mapping all their connections is an immensely difficult task, for which we currently do not have the imaging tools. However, multiple technologies have been and are being developed to tackle this problem.

Neuronal tracers are considered the gold standard for animal neuronal connectivity, as they trace a number of neuronal paths throughout the animal brain [7], [8], with the mouse brain having been studied and mapped extensively [7], [9], [10]. However, the limited ability to study human brains, in addition to the experimental effort for the multiple injections (and animals) and imaging sessions needed to cover a subset of the neurons of a typical animal brain makes this unique method inconvenient for standard assessment of structural connectivity.

At the nanometer level, electron microscopy can image at very high resolutions, providing us with exquisite images of human and animal brain at the sub-cellular level, where even single myelin layers can be clearly distinguished [11]. However, electron microscopy typically needs extended sample preparation procedures, which can alter the sample microstructure. Also, even when used in 3D, electron microcopy typically images a small part of the brain at these very high resolutions, usually extending to much less than 1mm^3^.

Alternatively, fluorescence microscopy methods can reach sub-micrometer resolutions, and visualize fibers in extended brain regions [12], while tissue clearing can help image large specimens such as whole mouse or rat brains [13]. However, tissue clearing usually involves tissue distortion and is more difficult for larger and non-perfusion-fixed human tissues. Furthermore, the use of structure tensor analysis, typically accompanying these direct microscopy methods for deriving orientation information, is prone to artifacts due to structures other than axons and can be difficult to apply in dense white matter regions where intensity gradients are small.

Imaging methods that directly probe axon orientations overcome many of these issues. 3D Polarized Light Imaging (3D-PLI) exploits the birefringence of myelin to derive fiber orientations in tissue sections at micrometer resolution, with the possible field of view extending to the entire human brain [14]. However, it cannot recover crossing fibers within a pixel, though in-plane pixels are sometimes small enough to still resolve individual bundles crossing one another over several pixels (Zeineh *et al*. [15], Fig 4). Furthermore, the determination of the out-of-plane angle by 3D-PLI is challenging. Polarization-sensitive optical coherence tomography (PS-OCT) in serial, back-scattered mode also provides high-resolution orientation information based on the birefringence of myelin [16], [17] while facilitating image registration, but the method has otherwise similar limitations to 3D-PLI. Scattered light imaging (SLI) [18] partly overcomes those challenges, being able to resolve fiber crossings with micrometer resolution, while also providing information on out-of-plane fiber angles. However, SLI is a new method that still needs validation, especially regarding the quantification of the out-of-plane fiber angles.

Finally, the most commonly used method to provide brain-wide structural connectivity is diffusion magnetic resonance imaging (dMRI) [19], [20]. dMRI is sensitive to the anisotropic movement of water molecules, which is used as a proxy for local tissue orientation. However, resolution is typically limited to the hundred micrometers range *ex vivo*, while its signal is also sensitive to tissue microstructures other than myelinated axons, such as non-neuronal cells, extracellular matrix, intracellular components, etc. Most methods used for neuronal orientation analysis to validate dMRI are reviewed in Yendiki *et al*. [21]

X-ray scattering probes tissue microstructure by investigating tissue’s interactions with X-ray photons traversing the sample: photons elastically deviating from the main path are recorded by an area detector a few meters downstream. When photons probe a highly repetitive structure such as myelin, they constructively interfere at specific angles, forming Bragg peaks (hereafter referred to as “myelin peaks”). Moreover, this constructive interference happens at the plane formed between the photon beam and the direction of periodicity of the repeated structure. In the case of myelin layers, this plane is perpendicular to the nerve fiber orientation. This allows the use of the location of the peaks on the detector for determining the orientation of the myelinated axons (cf. Fig. 1 and Georgiadis et al., 2020 [22]). Use of neutrons instead of photons provides similar information, yet with lower resolution and specificity [23].

**Figure 1.**
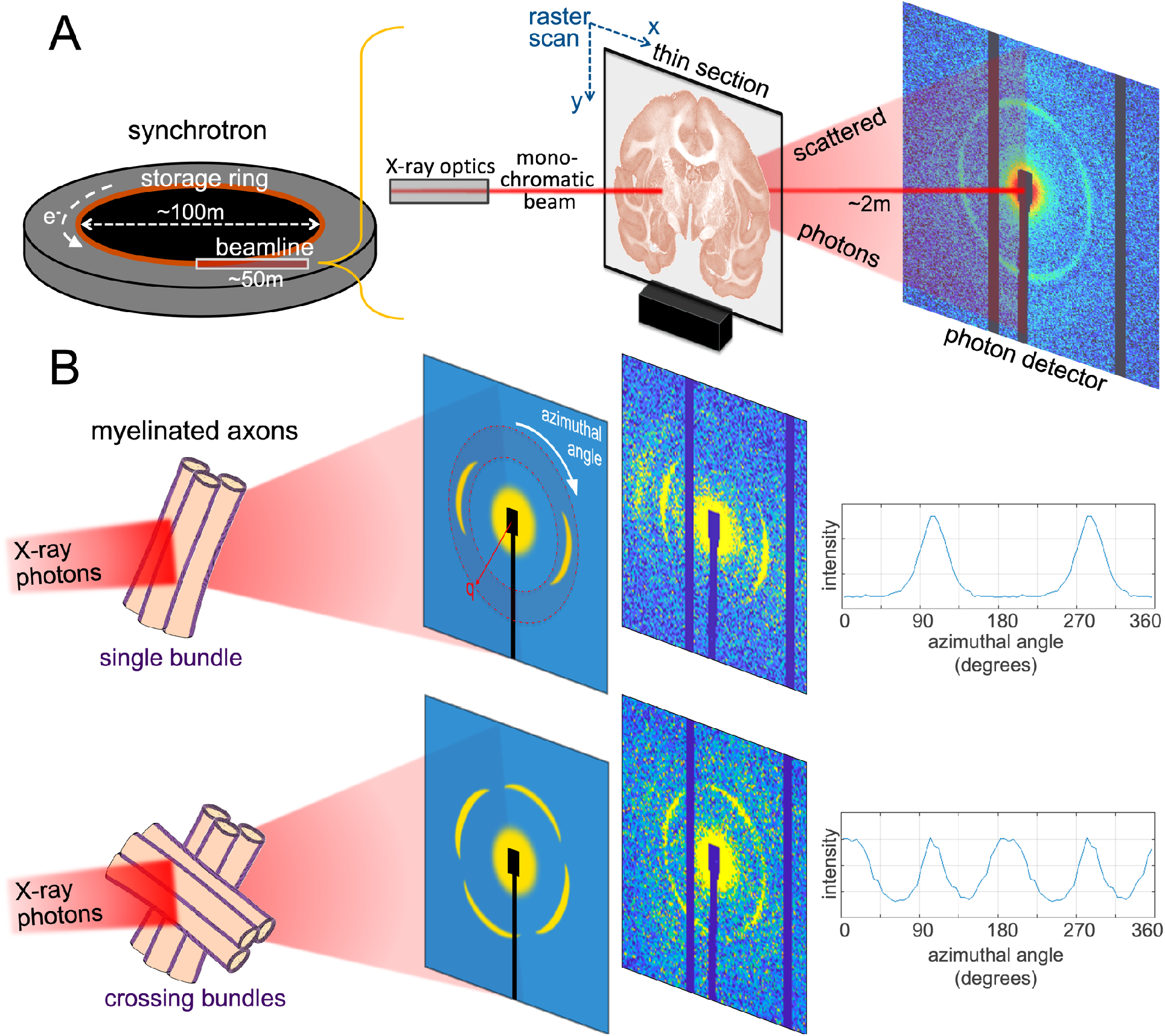
Detecting myelinated axon orientations using X-ray scattering. A) Schematic of experimental setup in the synchrotron beamline. The monochromatic X-ray beam impinges the thin section and photons are scattered at small angles according to their interactions with the sample’s nanostructure and captured by an area detector. B) Principle of detecting (crossing) orientations with sketch (left side) and real data (right side). The X-ray photons interact with the periodic myelin layers (left) and produce peaks in the resulting scattering pattern (middle). The radial position (distance from the pattern center, q) of the peak depends on the myelin layer periodicity d (q=2π/d), while the azimuthal position depends on the axonal orientations (with photons being scattered at a plane perpendicular to the axon orientation, cf. Georgiadis et al., 2020 [22]). To extract exact axonal orientation, the azimuthal profile of the myelin signal across a ring (in faint red in the top scattering pattern sketch) is plotted (right). The peaks are subsequently identified using the SLIX software (Reuter and Menzel, 2021 [30]). The position of the peaks in the x-axis (which are always 180° apart due to the center-symmetry of the pattern) reflects the fiber orientation angle. In the scattering patterns, the center (where the direct beam lands on the detector) is covered by a beamstop (usually including a photodiode), while the real scattering patterns (right) also have dark stripes (here in the up-down direction) corresponding to detector gaps that accommodate detector electronics. Also, in the real scattering patterns, multiple orders of the myelin peak are visible, albeit at much lower intensities, as expected by the Bragg law combined with the form factor of the myelin layer.

While determining 3D fiber orientation from scanning **small-angle X-ray scattering** (SAXS) signal has been shown in both thin sections using 3D scanning SAXS [22], [24], [25] and intact tissue specimens using SAXS tensor tomography [26]–[28], the use of scanning SAXS to disentangle crossing fibers in the animal and human brain has not yet been shown. Given the high importance of uncovering interwoven fiber trajectories in connectomics, we sought to determine whether SAXS can delineate crossing fibers. We first demonstrate the use of scanning SAXS to image in-plane crossing fibers in artificial crossings created using strips of human corpus callosum stacked on one another at different angles. In these configurations, we show that SAXS can detect crossing angles down to 25° and distinguish at least 3 crossing fibers. We also demonstrate the detection of crossing fibers in the white and gray matter of brains of multiple species that are actively investigated in connectomics, including mouse, pig, vervet monkey and humans. Taken together, we show scanning SAXS as an imaging tool that is both specific to axonal orientations and sensitive to crossing fiber detection. Scanning SAXS can uncover complex fiber geometries in challenging regions of animal and human brains and therefore serve as validation for state-of-the-art fiber orientation methods such as dMRI or SLI.

## 2 Methods

### 2.1 Brain samples

For creating the artificial crossings of fiber bundles, several sections from a human corpus callosum were used. The brain of a 66-y-o female donor with no sign of neurologic disease was obtained from the Stanford Alzheimer’s Disease Research Center Biobank (IRB-approved protocol #33727, including a written informed brain donation consent). It was cut in 1cm coronal slabs, immersion-fixed in 4% paraformaldehyde (PFA) for ∼2 weeks, and subsequently stored in a solution of 1x PBS + 0.02% sodium azide to deter bacteria growth. The body of the corpus callosum was excised using a scalpel, and 60μm sections were cut using a vibratome (VT1000S, Leica Micro-systems, Germany). The corpus callosum from several consecutive sections was manually cut into small, parallel strips of a few millimeters’ width. The tissue strips were manually stacked, aiming to create geometries of two or three fibers crossing, similar to the validation used for SLI [29]. For scanning, the stacked white matter strips were hydrated by phosphate-buffered saline (PBS) and placed between two 150μm-thick (#1) cover slips. Cover slips were glued around the edges (Super Glue, The Gorilla Glue Company, USA) to avoid sample dehydration.

The mouse brain sections investigated in this study were previously analyzed in [22] to derive single nerve fiber orientations for each tissue pixel. In brief, a vibratome was used to obtain one 25μm- and one 50μm-thick coronal section from a 5-month-old C57BL/6 female mouse brain, which had previously been fixed by transcardial perfusion with PBS and 4% PFA and then immersed in 4% PFA at 4° C for 48hrs. Procedures were within the animal license ZH242/14 of the Animal Imaging Center of ETH Zurich/University of Zurich. Sections were stored in PBS at 4° C. For scanning, they were enclosed in a PBS bath, within two thin Kapton films (Benetec GmbH, Switzerland) and a custom-made metal frame (see Suppl. Fig. 1 in [22]).

The pig brain was extracted from a female 10-week-old micro-Yucatan minipig, following institutional approval. The brain was immersion-fixed in 4% PFA for a month. A coronal slab was cut out of the left hemisphere, and a vibratome (VT1000S, Leica Microsystems, Germany) was used to cut 100-μm sections. The section together with a minute amount of PBS was placed between #1 cover slips, which were then glued around the edges to avoid sample drying.

The vervet monkey brain section was obtained as described in Menzel *et al*. [29] (IACUC #A11-219). Briefly, the extracted brain was fixed in 4% PFA and embedded in a 20% glycerin and 2% dimethyl sulfoxide solution before freezing. A cryotome (*Polycut CM 3500*, Leica Microsystems, Germany) was used to obtain 60μm-thick sections, stored in 20% glycerin. For the scanning SAXS experiments, one section was immersed in PBS for a few weeks, and then placed between #1 coverslips similar to the pig section and corpus callosum strips.

Finally, the human hippocampus section was obtained from the Stanford ADRC Biobank, from the brain of a 80-y-o female donor with a pathologic diagnosis of low Alzheimer’s disease (AD staging A1B1C2, Amyloid 1, Braak’s 1, Cerad 2, also diagnosed with hippocampal sclerosis of aging, with positive TDP-43 staining). After extraction, the brain was fixed with 4% formaldehyde, cut into 1cm coronal slabs, and stored in PBS. The left hippocampus was excised and a 75μm section was obtained using a vibratome (VT1000S, Leica Microsystems, Germany). For scanning, similar to the mouse sections, the human brain section was immersed in PBS and enclosed within two thin Kapton films, surrounded by a metal frame.

All animal procedures were in accordance with the National Institutes of Health guidelines for the use and care of laboratory animals and in compliance with the ARRIVE guidelines.

### 2.2 SAXS scanning

Each section was raster-scanned by the X-ray beam (details in Table 1). SAXS data were collected by a Pilatus 1M detector (Dectris AG, Switzerland) ∼2 meters downstream. A Pilatus 2M was used for the mouse data. The beam diameter and motor step sizes were matched (cf Table 1), constituting the effective pixel size for each scan.

**Table 1.** Experimental details for SAXS scans

|  | 2x/3x crossings | mouse | pig | vervet | human |
| --- | --- | --- | --- | --- | --- |
| Beamline, Synchrotron* | 4-2, SSRL | cSAXS, SLS | 4-2, SSRL | 4-2, SSRL | 4-2, SSRL |
| Container | #1 coverslips | Kapton tapes | #1 coverslips | #1 coverslips | Kapton tapes |
| Section thickness ($\mu\text{m}$) | 2×60, 3×60 | 25 \| 50 | 100 | 60 | 75 |
| Beam energy (KeV) | 15 | 12.4 \| 16.3 | 15 | 17 | 11 |
| Beam diameter ($\mu\text{m}^2$) | 125 | 25 \| 50 | 100 | 125 | 100 |
| Motor step size ($\mu\text{m}$) | 125 | 25 \| 50 | 100 | 125 | 100 |
| Field of view (x-y matrix) | 38 × 39 \| 42 × 46 | 256 × 400 \| 80 × 102 | 140 × 199 | 309 × 480 | 136 × 250 |
| Exposure time/frame (ms) | 500 | 50 \| 300 | 500 | 600 | 150 |

\* SSRL: Stanford Synchrotron Radiation Lightsource, SLS: Swiss Light Source

### 2.3 SAXS data analysis

A silver behenate (AgBe) standard with known periodicity (5.84nm) was used to identify the scattering pattern center and calibrate the sample-to-detector distance and inverse space units. Each scattering pattern was then segregated into 72 azimuthal segments (5° steps). The signal intensity modulation along the radial direction was obtained for each segment. Photon counts from pixels in 180-degrees opposite segments were averaged due to the center-symmetry of the pattern. This also allows filling in missing information from the regions that correspond to detector electronics (dark bands in scattering patterns, cf. Fig. 1).

The myelin-specific signal for each segment was isolated using procedures similar to those described in Georgiadis *et al*. (2021) [28], i.e. by fitting a polynomial to the radial signal intensity curve, and isolating the 2^nd^ order myelin peak (within the red ring in Fig. 1), which is the most prominent peak in myelin scattering patterns [28]. This allowed creating azimuthal profiles for each point, plotted in Fig. 1B, right. These reflect the orientation of the different populations of myelinated fibers within each probed volume (pixel size × section thickness).

To quantify and visualize the orientations of crossing nerve fibers, by detecting the peaks and identifying their position, the SLIX software [30] available on https://github.com/3d-pli/SLIX was used. This software has been developed to detect up to three different fiber orientations per pixel for scattered light imaging [18], where photons also scatter off the sample depending on nano/microstructure orientation and generate angle-dependent azimuthal profiles and associated peaks [31], a nearly identical computational problem to that with SAXS. Feeding the SAXS azimuthal profile data from each tissue pixel to SLIX produced the single or crossing in-plane fiber orientations per pixel. The SLIX *“--smoothing fourier*” parameters used were 0.4/0.225 (default settings suggested by [30]), and 0.1/0.01 for obtaining the main hippocampal fiber orientations (this smoothens the curve only preserving the main peaks). The length of the vectors in vector maps is weighted by the average myelin signal in each pixel.

In the orientation encoded colormaps (e.g., in Figs. 2-7A), each pixel is split into 4 quadrants, to accommodate 4 possible SLIX-derived fiber orientation colors. If there is a single orientation, all 4 quadrants have the same color. If there are 2 orientations, diagonal quadrants have the same color. If there are 3 orientations, 3 quadrants have corresponding color, and the 4^th^ is black.

**Figure 2.**
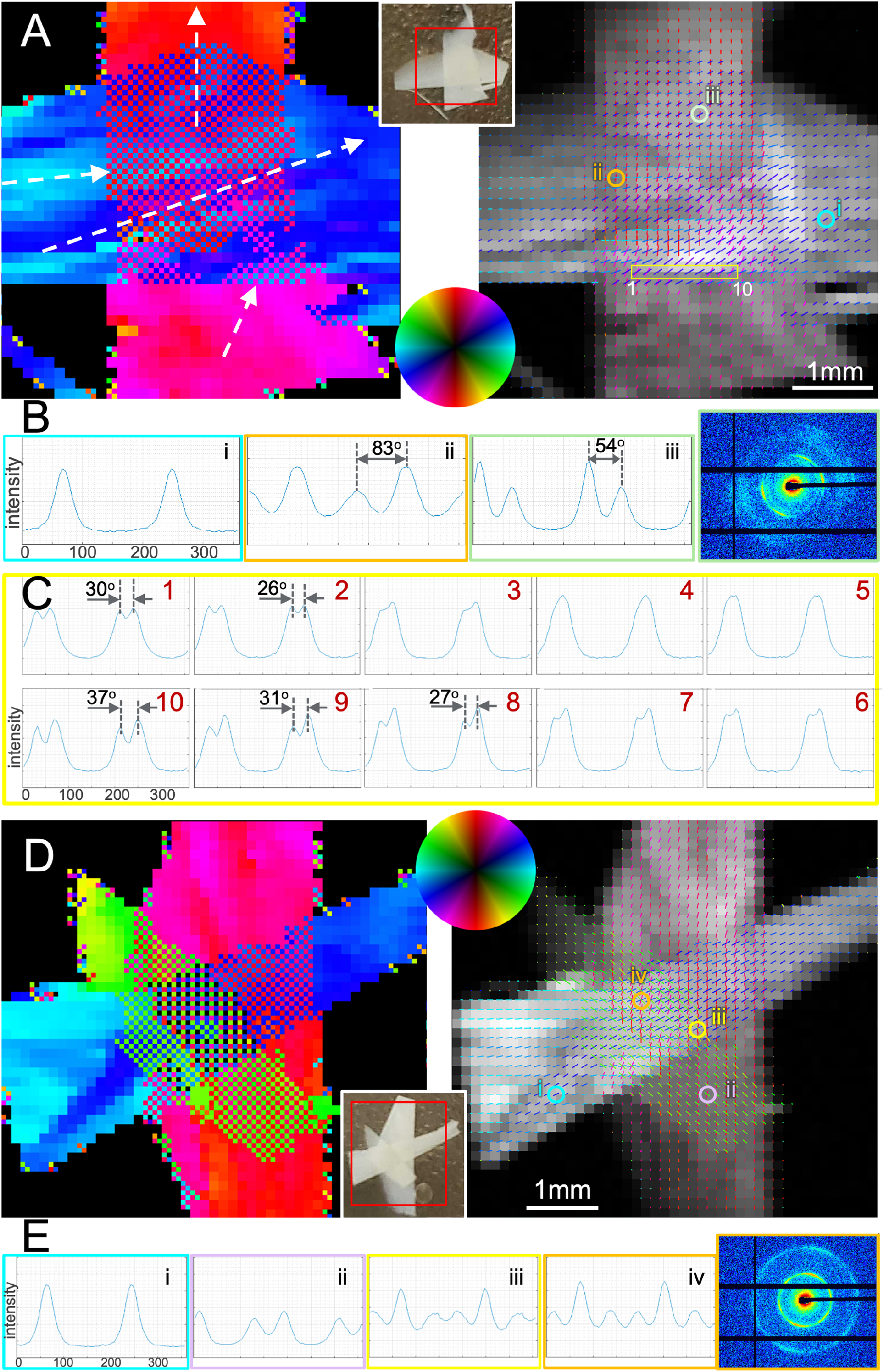
Imaging fiber orientations in artificial 2x and 3x crossings using strips of human corpus callosum. **A)** Two fiber bundles crossing, with fiber orientations for each pixel encoded in its color (left) or plotted as colored bars (right). Orientation is color-encoded according to the color wheel. Inset: Photo of the fiber strips within the coverslips, with scanned area in red rectangle. **B)** Azimuthal profiles (azimuthal scattering intensity across the myelin peak, cf. Fig. 1) for the circled points in (A). Image outline colors correspond to the colors of the circles in (A). **C)** Azimuthal profiles of 10 subsequent points surrounded by yellow rectangle in (A). Points show transition between two peaks (points 1-2), to one merged peak (points 3-7), and back to two peaks (points 8-10). Limit of detection using SLIX seems to be a crossing angle of approximately 25-30 degrees. **D)** Three fiber bundles crossing, with fiber orientations for each pixel encoded in its color (left) or plotted as colored bars (right). Orientation is color-encoded according to the color wheel. Inset: Photo of the fiber strips within the coverslips, with scanned area in red rectangle. **E)** Azimuthal profiles from select points in (D), with one (cyan), two (magenta), and three (orange & yellow) crossing fibers. As explained in the Methods, in the colored fiber orientation maps (A), (D), each pixel is split into 4 quadrants, to accommodate 4 possible SLIX-derived fiber orientation colors. If single orientation, all 4 quadrants have same color. If 2 orientations, diagonal quadrants have same color. If 3 orientation, 3 quadrants have corresponding color, 4^th^ is black.

### 2.4 3D-PLI

The determination of fiber orientations using 3D-PLI was done in an unstained section of a different C57BL/6 mouse brain. The mouse brain was immersion-fixed in 4% buffered PFA. After cryoprotection (20% glycerol), the brain was deep frozen at −70° C. The brain was serially sectioned along the coronal plane at 60 μm thickness using a cryotome (Leica Microsystems, Germany). Sections were mounted on a glass slide, embedded in 20% glycerol, cover-slipped and sealed with nail polish. The protocol was approved by the institutional animal welfare committee at the Research Centre Jülich, in accordance with European Union (National Institutes of Health) guidelines for the use and care of laboratory animals.

Polarimetric measurements of the section were done using the polarizing microscope LMP-1 (Taorad GmbH, Germany). The LMP-1 provides a field of view of 2.7 × 2.7mm^2^ and a pixel size of 1.3μm. Whole mouse brain section scans were carried out tile-wise using a movable specimen stage and a rotating polarizing filter. For each tile, a stack of 18 images was acquired at equidistant rotation angles (±10°) within the range of 0° to 170°. The measured intensity profile for an individual pixel across the stack of image tiles describes a sinusoidal curve that depends on the spatial orientation of fibers within this pixel. The physical description of the light intensity profile was derived from the Jones calculus for linear optics and represents the basis for orientation analysis, as detailed in [32].

## 3 Results

### 3.1 Validation with artificial crossings - SAXS detects up to three fibers down to 25° crossing angles

Analysis of the 2x-fiber configuration (Fig. 2A) shows that SAXS can recover crossing fiber populations. Strong fiber crossings are observed at most pixels in the overlapping region, with varying angles between the fibers in the two tissue strips. In general, the vertical strip shows fibers mostly running up-down (red color), with some rightward orientation of the bottom half (pinkish color). On the other hand, the horizontal strip seems to contain fibers running left-right with an inclination of ∼30° (blue color), e.g., point [i] in right part of Fig. 2A, plotted in Fig. 2B, with some parts having fibers running almost horizontally (cyan). In one such region, at the middle-left of the overlapping region, the fibers seem to cross at almost 90° (Fig. 2A/B, point [ii]). In the upper part of the overlap region, fibers from the two strips seems to cross at angles ∼50° to 60° (cf Fig. 2A/B, point [iii]). While most crossings are resolved, there are small regions (e.g., in the lower and right parts) where the angle between fibers decreases to the point where the two fiber populations are indistinguishable. This resolving limit is shown in the 10 subsequent points highlighted with the yellow rectangle in Fig. 2A, of which the azimuthal profiles are displayed in Fig. 2C. Starting from the pixel to the left (#1), the angle between crossing fibers decreases from 30° to 26° (#2), while in pixels #3-7 the two peaks are not distinguished by the algorithm. At pixels #8-10, the two separate fibers are detected again, with angles from 27° to 37° respectively.

The 3x-fiber configuration (Fig. 2D) showcases the ability of SAXS to detect at least three crossing fiber tracts within a pixel. The fiber orientations from the 3 tissue strips have an average of 60° between them, with the respective peaks clearly distinguishable, Fig. 2E. The majority of pixels in the region of overlap demonstrates 3 orientations, though in some regions only 2 crossing fibers are detected.

### 3.2 Mouse brain – detecting fiber crossings in myelinated areas

The mouse brain scanning SAXS data analysis using SLIX yielded a detailed fiber orientation map for the 25μm-thin section (Fig. 3A,B,E, the latter shows a zoomed-in view of the region of the box from B), revealing intricate crossings in myelinated brain areas. In the white matter, caudoputamen (CPu) fibers merge with the corpus callosum (cc), while some (white arrows) cross the lateral corpus callosum and the external capsule (ec) en route to the cortex, specifically the supplemental somatosensory area (SSs) (anatomic regions delineated on 3D-PLI in D). These crossings can also be inferred from the fiber tract trajectories seen using micrometer-resolution imaging of a section at a ∼300μm anterior plane from a different mouse with 3D-PLI (Fig. 3C,D, taking into account that 3D-PLI has limited sensitivity to crossing fibers within a pixel). They are more clearly visible in the axons seen by the tracer studies depicted in Fig. 3F-H, from the Allen Mouse Brain Connectivity Atlas [7] (https://connectivity.brain-map.org/), corresponding to experiments 297945448-SSs and 520728084-SSs, which include injections in the supplemental somatosensory area. These axons can be clearly seen crossing primarily the lateral side of the corpus callosum and the external capsule to reach the caudoputamen.

**Figure 3.**
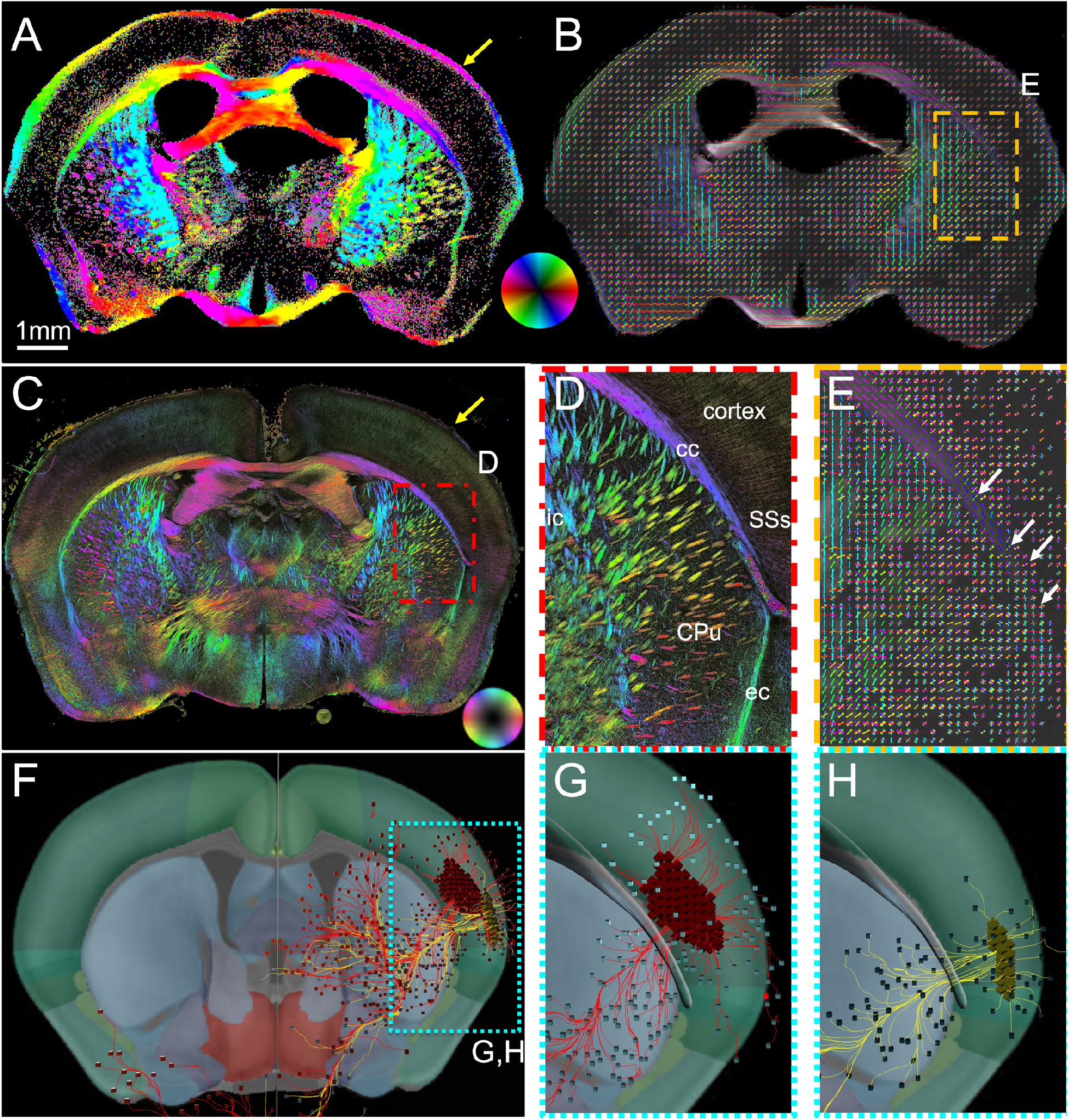
Fiber orientations in the mouse brain using scanning SAXS. **A**,**B)** Fiber orientations map of 25μm-thick section, with orientation color-encoded according to color wheel (A), or with orientation-encoded colored bars (B). To facilitate viewing of the image, one vector for each 5×5 pixel set is shown in (B). **C)** 3D-PLI image showing fiber orientations in a 60μm-thick mouse brain section, at a plane ∼300μm anterior compared to the section in (A-B). **D)** Zoom-in from (C). Yellow arrow points to molecular layer of cortex. **E)** Zoom-in of orange boxed region in (B), where fibers spreading from the internal capsule through the caudoputamen cross the corpus callosum and external capsule (white arrows) to continue radially into the cortex, and radial fibers reach the outermost molecular layer of the cortex and cross with the circumferential myelinated fibers along the brain surface (yellow arrow). **F-H)** Visualization of the Allen Mouse Brain Connectivity Atlas experiments 297945448-SSs and 266175461-SSs (https://connectivity.brain-map.org/projection/experiment/297945448 and https://connectivity.brain-map.org/projection/experiment/520728084), which included injections in the supplemental somatosensory area, showing axons in red and yellow colors respectively from a C57BL/6J mouse tagged with green fluorescent protein. **F)** Posterior view of both experiments, with a more anterior coronal section at the back for anatomical reference. **G-H)** Zoom-in from the cyan box in (F), with the corpus callosum in 3D rendering in gray color, rendered semi-transparent to enable visualization of cortical-caudoputamen fibers crossing it. Images created using the Atlas’ 3D viewer (https://connectivity.brain-map.org/3d-viewer). **cc**: corpus callosum, **ic**: internal capsule, **ec**: external capsule, **CPu**: caudoputamen, **SSs**: supplemental somatosensory area.

At the same time, crossings at the edge of the cortex, at the molecular layer, are also visible with scanning SAXS, where myelinated fibers running circumferential to the brain surface to connect to radial fibers from the rest of the cortex (yellow arrow in Fig. 3B), with the circumferentially oriented axons also visible in 3D-PLI (yellow arrow in Fig. 3C).

### 3.3 Pig brain – imaging complex fiber architectures in white matter

Contrary to the commonly studied lissencephalic rodent brain, the pig brain is gyrencephalic, meaning it has a folded structure containing gyri and sulci. This together with its bigger size results in a greater number of myelinated fibers with higher structural complexity of fiber tracts and multiple tract crossings (Fig. 4). The main regions of crossing fibers are enclosed in dotted boxes in Fig. 4B (labelled (i), (ii), and (iii)), and zoomed-in in Fig. 4C. The regions seem to be rich in 2x-fiber crossings, while some triple crossings can also be observed.

**Figure 4.**
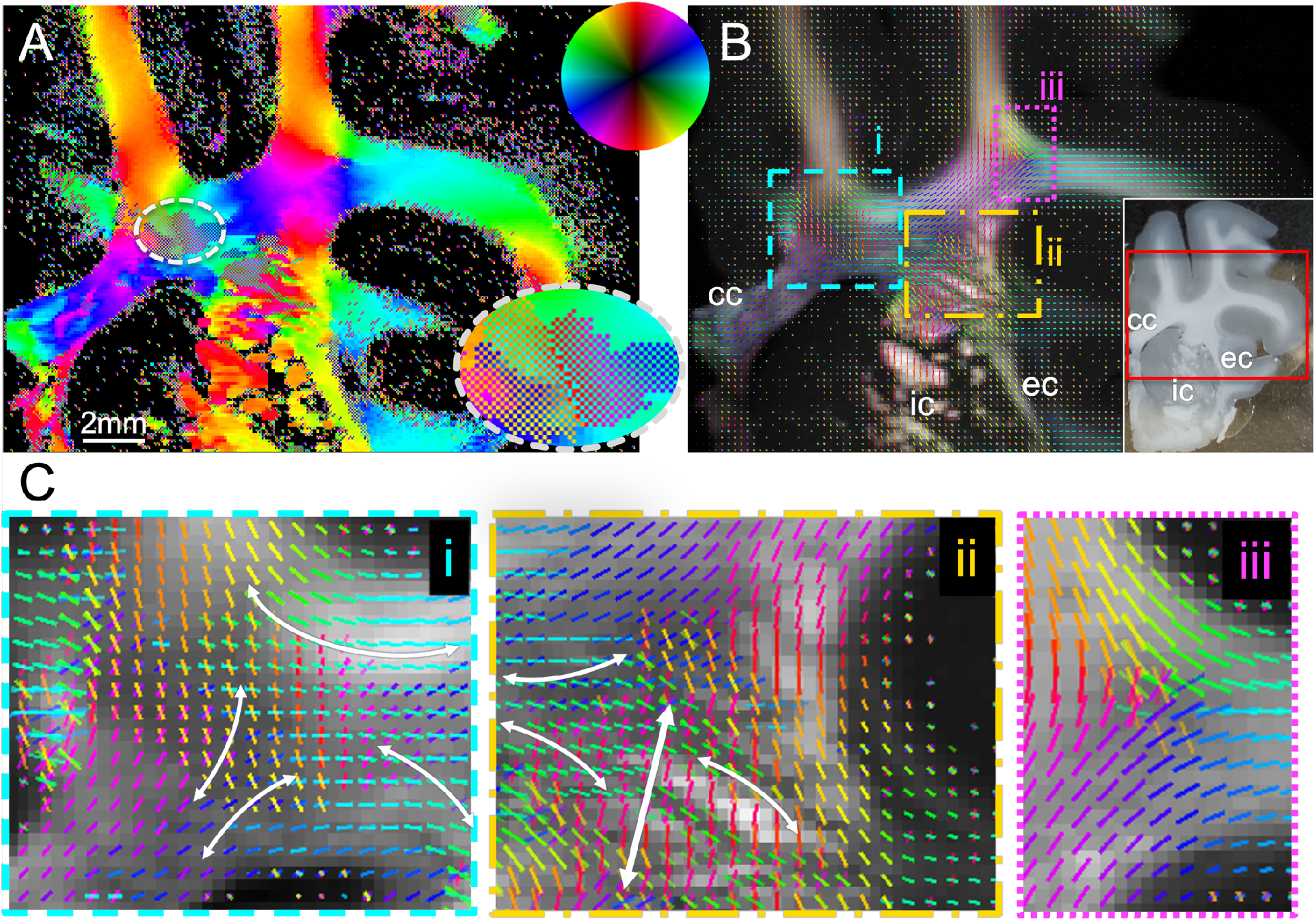
Fiber orientation analysis in the pig coronal section containing part of the corona radiata. **A)** Fiber orientation map of 100μm-thick section, with color encoded fiber orientations according to the color wheel. Inset shows zoomed-in region of crossing fibers, with the colors appearing in a checkerboard pattern, due to multiple orientations per pixel. **B)** Fiber orientations, represented by orientation-encoded colored bars for each 2×2 pixel set. The inset shows a picture of the pig hemisphere section mounted between the two coverslips, with the red rectangle indicating the region scanned. **C)** Subregions of the pig section, highlighting areas of various double and triple crossings. **cc**: corpus callosum, **ic**: internal capsule, **ec**: external capsule.

For instance, region (i) (cyan) contains callosal fibers from the lower left part that curve rightwards or upwards (bottom left pair of white lines), crossing with horizontally-traversing subcortical U-fibers (top arrow) and corticospinal fibers that go to the internal capsule (far-right arrow). Similarly, in region (ii) (orange), the fibers from the left side (including callosal and subcortical U) or from right side (external capsule) cross with the numerous almost vertical corticospinal fibers from the internal capsule (vertical thick white arrow). Some corticospinal fibers from the superior frontal gyrus (bottom right curved arrow in i) likely join this vertical portion of the internal capsule. Finally, in region (iii) (purple), another subcortical U-fiber population crosses with presumed callosal and internal capsule fibers.

### 3.4 Vervet brain – detecting crossings in challenging primate white matter geometries

The orientation analysis in the vervet brain section produced detailed maps of single and crossing fiber bundles (Fig. 5). Maps of the brain are seen in Figs 5A-B, showing fiber bundles in the corpus callosum (cc), corona radiata (cr), fornix (fx), caudate nucleus (cn), thalamus (th), internal capsule (ic), cerebral peduncle (cp), optic tract (ot), putamen (pt), external capsule (ec), extreme capsule (xc), and hippocampus (hp). The regions with the most prominent crossing fiber populations are those in the corona radiata, Figures 5C and D. There, at least two fiber populations cross, specifically the fibers radiating out of the corpus callosum (mostly left-right direction), and the fibers coming from the internal, external and extreme capsules (mostly superior-inferior direction).

**Figure 5.**
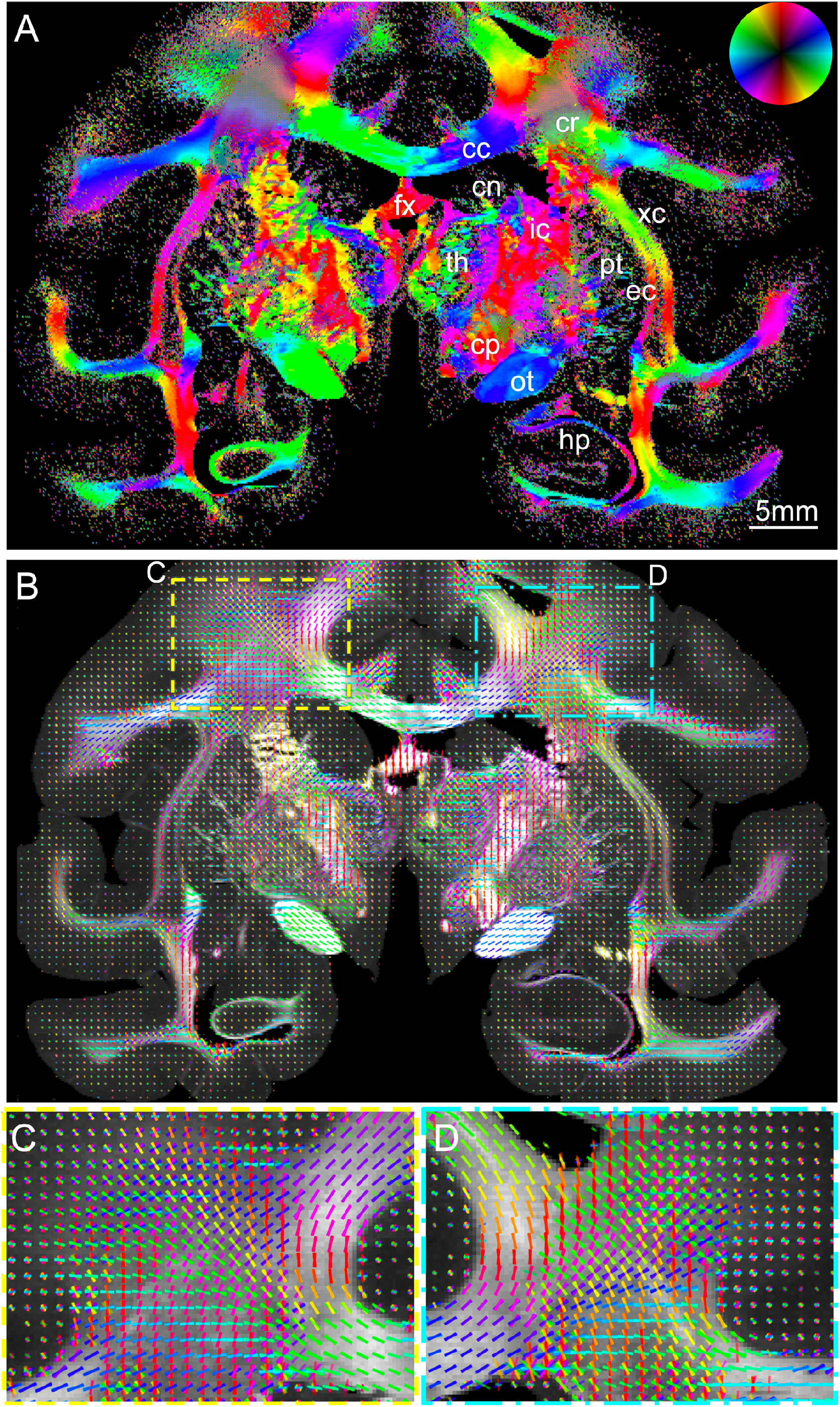
SAXS fiber orientation analysis in the vervet monkey brain. **A)** Fiber orientation map of 60μm-thick section (section nr. 518), with fiber orientations color-encoded according to the color wheel. **B)** Fiber orientations for the same section, represented by orientation-encoded colored bars for each 2×2 pixel set. **C-D**) Zoomed-in images of the left and right corona radiata from the vector map. **cc**: corpus callosum, **cr**: corona radiata, **fx**: fornix, **cn**: caudate nucleus, **th**: thalamus, **ic**: internal capsule, **cp**: cerebral peduncle, **ot**: optic tract, **pt**: putamen, **ec**: external capsule, **xc**: extreme capsule, **hp**: hippocampus

### 3.5 Pushing the limits: imaging gray matter crossings at the mouse cortex

The cortex is known to contain myelinated axons, with complex fiber architecture that can include crossings. Along these lines, apart from the crossings in primarily white matter regions shown in Figure 3 and also depicted here in Fig. 6C, multiple crossings were also detected in gray matter regions of the mouse brain. Figure 6D displays part of the mouse cortex, where myelinated fibers radiate from the corpus callosum towards the periphery in all parts of the cortex. In addition, fibers are observed running through the cortex in a direction tangential to the corpus callosum (e.g. Fig. 6D, in mostly left-right direction, dashed colored arrows show the radial axon orientations, depicted by color-encoded lines).

**Figure 6.**
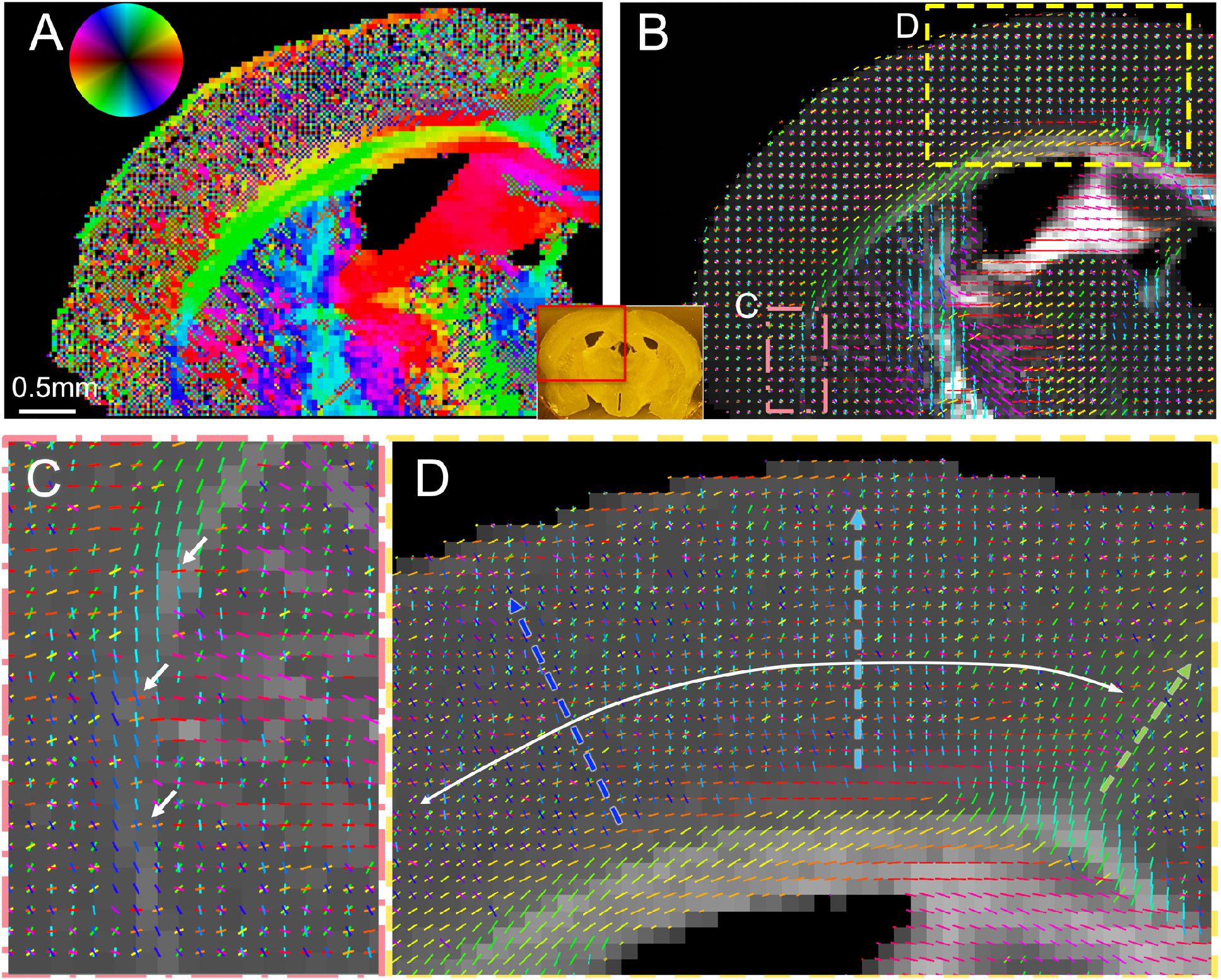
SAXS of crossing fibers in mouse cortex. **A-B)** Fiber orientation maps of 50μm-thick section, located ∼200μm anterior to section in Fig. 3 (section with scanned region in red rectangle shown in inset). (A) is color-encoded map, (B) is vector map. **C)** Zoom-in of orange-red boxed region in (B), showing white matter fibers radiating from the internal capsule through the caudoputamen and crossing the external capsule fibers (in regions of white arrows) towards the lateral cortex. **D)** Zoom-in of yellow boxed region in (B), including part of the cortex, the corpus callosum and the cingulum. Fibers radiating out of each region of the corpus callosum towards the cortex, are depicted by the blue, cyan and green arrows. Vectors in each pixel in C and D correspond to the nominal image resolution, namely 50μm.

### 3.6 Human hippocampus – detecting subtle secondary fiber tract crossings in human brain

The human hippocampus is a gray matter structure with numerous interwoven white matter pathways of immense complexity and importance to memory formation, which are altered in neurodegeneration and diseases such as Alzheimer’s disease and epilepsy. Analysis of the human hippocampus section provided detailed maps of myelinated axon orientations across hippocampal subfields and the various hippocampal fiber tracts, as demonstrated in Figure 7.

**Figure 7.**
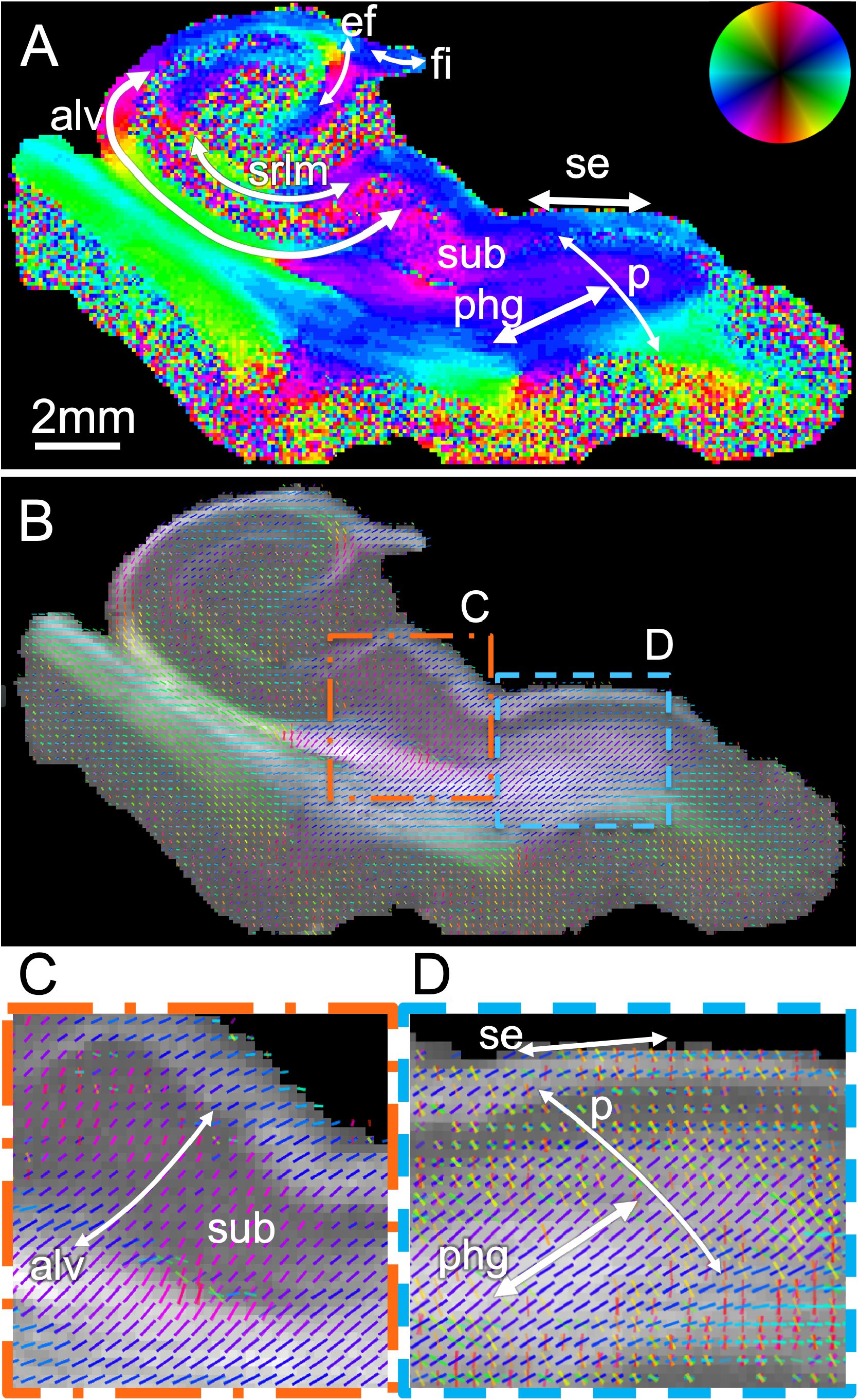
Fiber orientation analysis in the human hippocampus. **A)** Fiber orientation map of a 75μm section, with fiber orientations color-encoded according to the color wheel. **B)** Fiber orientations represented by orientation-encoded colored bars for each 2×2 pixel set. Only main orientation is depicted. **C)** Zoomed-in image of the orange box in (B), showing forniceal tract (fo) fibers running through subiculum. **D)** Zoomed-in image of the blue box in (B), depicting both primary and secondary fiber bundles in each pixel. **p**: perforant pathway, **phg**: parahipocampal gyral white matter, **se**: superficial entorhinal pathway, **sub**: subiculum, **fo**: forniceal path, **srlm**: stratum radiatum lacunosum and moleculare, **ef**: endfolial pathway, **fi**: fimbria.

Figure 7A shows the color-encoded main fiber orientation for each pixel (achieved by adjusting SLIX parameters to only detect the main peak in the line profiles, cf. Methods), revealing part of the complex anatomy of the hippocampus and the geometry of its fiber bundles. Primary fiber orientations are also depicted visually in Figure 7B, in both white and gray matter areas. The main white matter tracts can be seen there, including the alvear/forniceal (alv), perforant (p), stratum radiatum lacunosum and moleculare (srlm), endfolial (ef) and superficial entorhinal (se) pathways. Subicular fibers heading to/from the alveus (Fig. 7C) are also seen crossing the gray matter. In Figure 7D, the cyan box in (B) is shown in higher resolution, demonstrating secondary and tertiary orientations of myelinated axons within white matter pixels. Multiple fiber orientations can be seen in most white matter pixels, meaning that in the complex hippocampal anatomy and connectivity, each pixel rarely contains fibers from a single bundle. For instance, the perforant pathway (p) is seen crossing the angular bundle within the parahippocampal gyrus (phg) just below the subiculum (sub), and reaching the superficial entorhinal (se) pathway within the stratum radiatum lacunosum and moleculare (srlm) and crossing the subicular forniceal bundle (alv) from 7C.

## 4 Discussion

In this study, we demonstrate that SAXS can distinguish at least 3 distinct crossing fiber populations, at crossing angles as low as ∼25°, and we show crossings in white and gray matter across different species. This separation is uncovered by analyzing the peaks in the azimuthal profiles from each point’s scattering pattern, isolating the myelin signal corresponding to the periodicity of myelin. This work is a critical new direction to scanning SAXS capabilities shown previously, building upon the determination of main 3D fiber orientations in sections [22], [24], comparison to MRI, and quantification of myelin levels, orientation, and integrity in volumetric specimens [28].

### 4.1 Crossing angle detection limit

Given the experimental conditions and analysis tools presented in this study, the current limit of detection seems to be approximately 25° -30°. This resolving power depends on a few factors: first, the signal-to-noise ratio; if less photons impinge the sample, it is possible that noise in the azimuthal profiles could hamper peak detection. However, the current experiments with typical experimental conditions in scanning SAXS seem to yield signals with a high signal-to-noise ratio, which are expected to be robust to such effects. Secondly, there is an inherent resolving limit to the method, which depends on the “fiber response function”, i.e., the angular distribution of the signal coming from a single coherent, unidirectional fiber bundle. In theory, the SAXS signal from a single straight fiber would be exactly at 90 degrees to the fiber orientation and have an azimuthal profile with two peaks of minimal width. However, in practice myelinated axons have some curvature and undulations, and each pixel probed by the beam contains multiple fibers with a certain orientation distribution. Under these circumstances, the effective fiber response function has an azimuthal profile with a distribution similar to the peaks in Figures 1 or 2. Taking these considerations as well as data from the current experiment into account, we speculate that the crossing angle detection could be lower at approximately 20° ; possible improvements to the 25° -30° presented here could come from higher signal-to-noise ratio experiments (i.e. using higher flux) or even more sensitive peak detection algorithms, as discussed below. Finally, another case where crossing fibers can remain undetected is a scenario where the secondary fiber population contains considerably less myelinated axons (lower number of axons, or axons with a lower degree of myelination). In that case, its scattering peak might not be able to be detected, unless the signal-to-noise ratio is very high or a micrometer-resolution scan can visually resolve the myelinated axons contained in different tracts. Such resolution can be attained by decreasing beam diameter and motor step size, or by use of other orientation-sensitive microscopy methods, such as scattered light imaging [18], [29], [31].

### 4.2 Comparison to other methods

Scanning SAXS has different advantages and shortcomings compared to the multiple other methods employed towards detecting fiber orientations -and crossing fibers-in the animal and human brain. Since diffusion MRI is the only modality which can be applied both *in vivo* and *ex vivo* with minimal sample preparation, other methods aim to overcome its shortcomings.

Limitations of diffusion MRI mostly stem from the fact that it uses the directional movement of water molecules as a proxy for fiber orientation. As a result, i) it is difficult to translate findings from *ex vivo* to *in vivo*, since water content and motility are altered in fixed samples, ii) movement of water is also restricted or hindered by all structures in the brain [33], including membranes of other cells, organelles, extracellular matrix and vesicles, cell cytoskeleton, vesicle walls, while water can passively or actively move across membranes [34], iii) fiber response function is inherently broader compared to SAXS, because even in the case of perfectly aligned fiber population, all the above factors, as well as the movement of water in random directions (to the extent allowed) within the neuronal somata, axons, and in the extracellular space, contribute to an angular dispersion of the signal. In addition, dMRI resolution is typically restricted to hundreds of micrometers per voxel, potentially corresponding to thousands of fibers, further contributing to the angular dispersion. All these make interpretation of diffusion MRI output very challenging and stress the need for validation methods [21].

As described, the main limitation of dMRI is lack of specificity to myelinated axons. This can be provided by scanning SAXS, confocal/multi-photon microscopy (possibly combined with clearing), and polarization-based methods (3D-PLI/PS-OCT). The limitation of polarization-based methods is that they are performed on 2D samples (sections or surfaces). On the other hand, confocal/multi-photon microscopy without clearing is limited in depth, and different antibodies can yield different results, depending on the tissue origin, preparation etc., while clearing is labor and time-intensive, typically results in tissue deformation, and is challenging for large specimens. SAXS on the other hand does not depend on staining given its specificity to myelin nanostructure, and X-rays can penetrate thick specimens. Here, we are only reconstructing tissue slices using SAXS. Future implementations of SAXS tensor tomography with crossing fiber detection may provide specificity across a tissue volume.

The second feature where dMRI lacks is in resolution, which is limited to hundreds of micrometers. Methods such as 3D-PLI, SLI, and PS-OCT can reach micrometer resolutions, since each image pixel contains orientation information, with SLI also being able to detect crossings in each pixel, while 3D-PLI and PS-OCT depend on their high resolution for visually resolving crossings. Analysis of brightfield or fluorescence images using structure tensor can provide resolution of few to several tens of micrometers, depending on the kernel size used. Electron microscopy is the highest resolution method, reaching sub-nanometer resolutions, but the field of view is typically less than a millimeter, and sample preparation is extensive and usually tissue-distorting. Finally, scanning SAXS orientation analysis can reach sub-micrometer resolutions [35], [36], as it depends on the beamsize and motor step size used, and crossings can be detected for each pixel.

Another important advantage of scanning SAXS versus other MRI validation methods is its ability to provide quantitative 3D orientations in isotropic voxels, both tomographically and in sections. Brightfield methods are only 2D, while 3D fluorescence imaging has a lower through-plane compared to in-plane resolution, and structure tensor analysis is challenging in dense white matter regions where intensity gradients are low. 3D-PLI can also provide 3D orientation, but the out-of-plane angle can be ambiguous, while SLI yields information on out-of-plane angles, but quantification has not yet been achieved.

Finally, SAXS, in its tensor tomography form [26]–[28], is the only method together with MRI that can be performed on intact specimens, with most other methods being limited to sections. Tissue clearing also enables 3D imaging of specimens, but tissue is usually distorted by the clearing process, and samples larger than few millimeters pose challenges in clearing, antibody penetration and imaging.

### 4.3 Crossings in white and gray matter from multiple species

Crossings in the brains of various species are shown in this study. The mouse brain scan revealed crossings in both white and gray matter. White matter crossings were seen mostly by cortical fibers and crossing the corpus callosum and external capsule on their way to the caudoputamen. The relative scarcity of white matter crossings is expected given the rodent brain relative simplicity compared to gyrencephalic brains. Nevertheless, X-ray scattering was also able to distinguish crossings of myelinated axons in gray matter regions, including tangential fibers crossing the more abundant radial ones [37], only previously reported using tracer studies [7], special microscopy methods with very high resolution [38], or diffusion MRI specially utilizing very high angular and spatial resolution [37], [39].

Fiber crossings in a Yucatan micropig brain are also reported. Since the pig brain is increasingly used in neuroscientific studies [40], including studies of white matter and structural connectivity [41], [42], it is essential that the white matter fiber orientations are studied and accurately mapped. In addition, the swine model is often used in biomechanical studies [43], where axonal orientation seems to play a role for axonal injury and for computational determination of local mechanical strains [44]. Biomechanical implications of crossing fibers have not been adequately studied to date, so combined imaging and mechanical experiments could help elucidate the structure-biomechanics relationship.

The primate (vervet monkey) brain exhibits similar architecture to the human brain, which makes it highly suitable for neuroscientific studies [45] of structure [46] and function [47]. Besides, most knowledge of gold-standard human connectivity comes from tracer studies in primates [8], [48], [49]. 3D-PLI of vervet monkey brain regions has provided micrometer resolution fiber orientations maps [46], while SLI has additionally enabled discerning crossing nerve fibers [18], [29], [31]. Our scanning SAXS maps provide detailed fiber crossings across the vervet monkey brain section. The corona radiata SAXS maps appear similar to corona radiata SLI maps [18], [31]. Combining the specificity to myelinated axons of SAXS with the high-resolution capabilities of SLI can potentially help to solve long-standing fiber orientation and tractography issues in brain connectivity, such as the high false-positivity rates of tractography algorithms [50], [51], or the challenges posed in complex fiber geometry regions where multiple fiber bundles combine and have in/through-plane trajectories [51], [52].

The human hippocampus is critical for new memory formation, and shows degraded function in neurodegenerative diseases such as Alzheimer’s disease (AD) [53]. However, its location towards the lower part of the brain, next to the mastoid air cells, its complex anatomy [54] and connectivity [55] as well as its relatively small size, makes it challenging to study with MRI [56], which is the most commonly used *in vivo* brain imaging modality. Despite that, the significance of the hippocampus has led to multiple approaches to study its detailed anatomy and connectivity. *In vivo* MRI studies have been able to segment the hippocampal subfields [57], and show the main and even microscopic fiber pathways [58], [59]. *Ex vivo*, high-resolution MRI has been commonly used to study hippocampal connectivity in excised specimens; early high-resolution anatomical and diffusion tensor imaging scanning could delineate the different laminae [60], while modern scanners and approaches combining scanning with histology allow detailed subfield segmentation and studies in the context of neurodegenerative diseases [61], [62]. In addition, micrometer-resolution scanning using polarized light imaging has allowed unprecedented insights into the hippocampal connectivity [15]. Here, we have shown myelinated-axon-specific maps of fiber orientations using X-ray scattering in a human hippocampal specimen, including dominant and secondary fiber pathways in white and gray matter. Such information can potentially reveal a loss of normal white matter pathways in neurodegenerative diseases, where demyelination has been observed, as well as loss of specific pathways such as the perforant pathways in AD. This combined with increasing data at the histopathologic [63], cytomolecular [64], as well as transcriptomic [65], [66] levels can investigate how myelin and oligodendrocytes might be central in the disease mechanisms.

### 4.4 Limitations

The study has a number of limitations. First, the ground truth for fiber orientations is an ongoing challenge. We attempted to overcome this by i) validating against strips of unidirectional fibers artificially superimposed to mimic pixels with crossings, noting that scanning SAXS has been proven to work in brain tissue for primary orientation [22], [28], and ii) comparing to a micrometer-resolution 3D-PLI image of a similar region of a mouse brain. However, even in the artificial crossings using corpus callosum strips, fibers are not fully aligned within each strip, so ground-truth crossing angles cannot be exactly determined. On the other hand, SAXS yields directional information from myelinated axons directly and with specificity, so the fiber orientations are relatively easily and directly interpreted compared to structural imaging or diffusion MRI, where myelinated-axon-specific orientation analyses include more complex algorithms and more assumptions.

SAXS scans can achieve a moderately high resolution while discerning crossing fibers; the highest resolution demonstrated here was in the first mouse brain section, reaching 25 micrometers, while resolution was up to 125 micrometers for the vervet brain, much lower than the resolution typically reached in 3D-PLI or SLI. Although SAXS scanning can be performed at very high resolutions, down to nanometer levels [36], practical considerations (beamline capabilities regarding beamsize, and time needed for raster-scanning an extended field of view at such resolutions) limit the resolution to typically tens of micrometers. This is below or at the same order as that of diffusion MRI, with the advantage of specificity to the myelinated fibers, making it thus a very good validation tool for similarly-sized samples.

Sensitivity to peak detection in the azimuthal profiles presents a challenge for future improvements. In this study, the SLIX software [30] used the SAXS azimuthal profile data as input, since in SLI photons also scatter anisotropically off material depending on structure orientation, and in both methods the position of the peaks in azimuthal profiles reveals the in-plane orientation of the fibers. However, by looking at the azimuthal profiles of pixels 3, 6, and 7 in Fig. 2, one can suggest that there are two peaks that are at an angle of ∼20° apart. This could indicate that a more sensitive peak-fitting algorithm, the subject of our future work, could possibly be able to tell these peaks apart, and thus resolve even lower angle crossings.

When it comes to tissue preparation for imaging, the presented scanning SAXS experiments were performed on thin sections. Although this is a very common approach used in histology or methods such as 3D-PLI or SLI, in many cases it is not desired, or not feasible. However, with the advent of SAXS tensor tomography [26], [28], such experiments can also be performed tomographically on whole specimens without sectioning. The challenge of computing both the tensor-tomographic reconstruction and depicting multiple fiber orientations per voxel will also be the subject of future investigations. This would provide a tomographic gold-standard in the axonal orientation field and enable a head-to-head validation of diffusion MRI orientation information on the same specimens.

Access to scanning SAXS is also a limitation because the required photon flux of micro-focused beam is currently only obtainable from synchrotrons. However, several sites worldwide provide appropriate beamlines for collaborative use. Continued improvement of instrumentation and analysis will enable higher sample throughput as well as scanning SAXS experiments on lab SAXS setups.

## 5 Conclusion

Accurate and specific imaging of crossing nerve fibers is an important challenge in neuroscience. In the experiments presented here, we use scanning small-angle X-ray scattering (SAXS) to show that detecting crossing fibers is feasible for at least three crossing fiber bundles and crossing angles down to approximately 25°, and applied the method across species, in mouse, pig, vervet monkey, and human brain, in gray and white matter. Overall, as scanning SAXS can provide specificity to myelinated axonal orientations, which are responsible for long-distance signal transmission in the brain, it has the potential to become a reference method for accurate fiber orientation mapping. Combination of scanning SAXS with micrometer-resolution imaging approaches, such as 3D-PLI or SLI, taking advantage of SAXS’s specificity and 3D-PLI/SLI resolution, could provide ground truth information on fiber orientations, yield accurate structural connectivity maps and be the basis for validation of diffusion MRI signals.

## Acknowledgments

We thank Samuel Baker from Stanford Comparative Medicine, Aileen Schroeter and Markus Rudin from ETH Zurich, Roger Woods from the UCLA Brain Research Institute and Donald Born from Stanford Neuropathology for providing the pig, mouse, vervet and human brain samples respectively, as well as all animal and human donors. The present work was supported by the National Institutes of Health (NIH), award numbers R01NS088040, P41EB017183, R01AG061120-01, R01MH092311, and 5P40OD010965, and by the European Union’s Horizon 2020 Research and Innovation Programme under Grant Agreement No. 945539 (“Human Brain Project” SGA3). 3D-PLI images were generated by utilizing the computing time granted through JARA-HPC on the supercomputer JURECA at Forschungszentrum Juelich (FZJ), Germany. SSRL, SLAC National Accelerator Laboratory, is supported by the U.S. Department of Energy, under Contract No. DE-AC02-76SF00515.The SSRL Structural Molecular Biology Program is supported by the DOE Office of Biological and Environmental Research, and by the National Institutes of Health, National Institute of General Medical Sciences (P30GM133894). The Pilatus detector at beamline 4-2 at SSRL was funded under National Institutes of Health Grant S10OD021512. The experiment on mouse brain sections were carried out at the cSAXS beamline, Swiss Light Source, Paul Scherrer institute, Switzerland, and the artificial crossings, pig, vervet and human sections in the 4-2 beamline of Stanford Synchrotron Radiation Lightsource, SLAC National Laboratory, USA.

## Data statement

SAXS data for all samples are deposited at: https://doi.org/10.5281/zenodo.7228131

The authors declare no competing interest.

